# Growth control as a central regulator for tuning the cellular context

**DOI:** 10.64898/2025.12.19.695408

**Authors:** Ángeles Hueso-Gil, Jesús Miró-Bueno, Ángel Goñi-Moreno

## Abstract

The cellular context interacts with genetic circuits, decisively defining their performance. However, contextual dependencies (the interplay between the host and the circuit) are often difficult to engineer rationally, leading to a lack of control over circuit behaviour. To address this challenge, we replaced the native regulatory machinery of the RNA polymerase (RNAP), the cell’s core protein-making machine, from the bacteria *Pseudomonas putida* KT2440 with an inducible system, enabling tunable growth regulation and thereby gaining systemic control over the entire cellular machinery. Specifically, this was achieved by placing key components, the *β* and *β*’ subunits of the enzyme, under the control of the XylS-Pm inducible system. By using its cognate chemical inducer, 3-methylbenzoate, cell’s growth can be controlled at will, enabling the precise tuning of the cellular context into distinct, stable states. We correlated genetic circuit behaviour with the cell’s growth state by observing the constitutive expression of a reporter gene and the performance of a collection of genetic NOT logic gates. Our results show that the modulation of contextual dependencies is specific to the circuit components but not a random process. A mathematical model allowed us to classify that modulation into three different categories for our library of NOT gates, based on the prediction of how RNAP availability shapes host-circuit interaction. Finally, using growth control as an input for a 2-input circuit lead to a NAND gate with the potential for morphological computing, where the cell’s physical body itself undertakes part of the information processing. Our findings indicate that growth control can be used as an engineering parameter, allowing us to search for optimal scenarios that enhance the potential of genetic tools.

## 1 Introduction

Engineering genetic circuits in living cells [1, 2] to perform predefined computations [3, 4] is now routine. However, this process is not (fully) predictable [5] yet, mainly due to the lack of a deep understanding of the interactions between native cytoplasmic components (i.e., the host context) and the synthetic devices. Unknown interactions, termed here as *contextual dependencies*, between the endogenous and the orthologous elements lead to an unpredictable performance of genetic circuits [6]. Despite great progress on working towards that predictability by designing circuit parts as interchangeable between species as possible [7], or via software tools that predict sequence performance [8], orthogonality has never fully been reached nor dependencies entirely understood. This highlights the fact that genetic circuits are far from electronic counterparts due to the inherent complexity of living systems [9]. Dealing with contextual dependencies is then an overarching challenge that needs further attention, position that underpins this work.

Controlling bacterial growth without encountering unexpected effects on the circuit of interest represents a current challenge of the engineering of biology [10]. Cell division and growth arrest has revealed as a tool with many possible applications but any considered strategy to gain control over this growth affects other cellular functions that should keep their independent balance, as bacteria decide where to locate their resources. One of the most well-known advantages of decoupling growth from the function of interest appears in bioproduction of compounds [11]. Strategies followed in pursue of this independent control are diverse. Many of them have been used for regulation of strain growth in microbial artificial consortia, being the most widely spread those that make use of antibiotic resistances, quorum sensing systems [12], co-dependent syntrophies [13], changes on media carbon sources [14], or depletion of nutrients such as nitrogen or phosphates [15]. However, all of these methodologies rely on media composition variations which sometimes show incompatibilities with further steps of the workflow or induce some stress responses for the bacterial population. On the other side, syntrophic co-cultures limit the flexibility of consortia strain ratios, as each strain reaches its point of stability at expenses of the limiting compound produced by the other. When ratios need to be tuned at desired and changeable proportions, other strategies should be followed.

Manipulation of transcription or translation rates by means of controlling ribosomes or the RNA polymerase (RNAP) respectively entails another way for bacterial growth control. In the microbial model *Escherichia coli*, it has been noticed that the number of ribosomes exposes the greatest constraint for resource restriction and directs the activity of the RNAP to *rrn* operons, [16], therefore some examples have tried to tune growth by means of ribosome control [17]. However, change in ribosome amount can lead to the rise of the alarmone ppGpp, which triggers astringent response, provoking a different physiological status of the cell that may not be suitable for certain processes [18]. Other studies have shown that not only translation levels but the ratio between the RNAP and ribosomes determines growth [19]. Focusing on the transcription and the RNAP, previous works have induced RNAP production of *E. coli* by the LacI-pTac expression system, gaining control over *E. coli* ‘s growth [20] [21]. Later works have inhibited the polymerase with an RNAP specific repressor, Gp2, inducing growth arrest in order to improve production rates [22]. In *Caulobacter* sp., growth phase has been changed by manipulating the ppGpp alarmone concentrations through CtrA due to *β* RNAP subunit is sensitive to its presence [23]. Not only native but also orthogonal RNA polymerases can also be used in combination to the native system with the aim to control two-strain consortia. Activating T7 RNAP for cloramphenicol resistance gene transcription using blue light, the balance of two *E. coli* populations could be manipulated at desired rates [24].

Despite growth control can clearly be reached by regulating general transcription levels of the bacteria, it is expected that this have a clear impact over physiological state of the cell that induce changes in the behavior of simple and more complex circuits. Some previous works have taken a look to the growth effects over heterologous gene expression and genetic circuits, revealing a very complex scenario. For example, it was studied that gene expression (gratuitous or not for the overall fitness) and growth are bidirectionally related at the level of translation: the correlation of mRNA with protein ratio changes with growth rate, specially due to changes in nutrients of the media [16]. Changes in growth rate not only affect to gene expression values but also to the diversification of bacterial strategies influencing over noise in that expression and bet-hedging. Some models have described that bacteria disposing on higher concentrations of carbon sources grow faster but entering in a more inefficient metabolism that translates into differences in fitness and noise, [25, 26], which often translates to variations in noise that determine cellular fate [27, 28]. Considering more complex implemented functions than the mere production of a single gene, it was determined that a positive feedback loop entangled with a growth modulator produces a bistability in growth, and, what is more intuitive, growth can change the functionality of a that circuit, so the interdependence between the circuit and the growth status happens clearly in both directions [29]. Another interesting observation pointed that the induction of growth can change the sensitivity of a citcuit to its inducer depending on the mechanism used for this growth manipulation, being more sensitive cells that are under growth arrest compared to those in exponential phase [30]. *In silico* models have also study the impact of growth defects over circuits. Including cellular resources such as ribosomes and RBS’s, bacteria under slow growth allow heterologous gene expression up to a limit that overburdened cells by high growth cannot reach, causing heterologous gene expression impairment in a complex net of interactions with other factors [31].

In this work, we have placed the rpoBC operon coding for RNAP *β* and *β*’ subunits under the control of XylS-Pm inducible system, using the host *Pseudomonas putida* KT2440. By inducing with the chemical 3-methylbenzoate (3MBz), we managed to control the growth of the developed strain from near total growth impairment to the normal *wt* rate. In order to understand how this manipulation affects to implemented functions, we have tested the influence of the external regulation of RNAP production over the expression of a constitutive GFP and 17 different NOT gates beared by low copy number plasmids. GFP constitutive expression showed how growth rates can be modified affecting not only to general expression levels but also to noise and reproducibility of fluorescence averages. On the other side, the examination of NOT gates behaviour under different growth regimes allowed to find RNAP requirement patterns for each circuit tested. A predictive mathematical model based on RNAP availability helped to classify these responses and uncovered unexpected emergent behaviors in some circuits. Therefore, a tunable growth strain was revealed to be not only a suitable chasis for diverse applications, but also a powerful tool for the study of the growth status of the cell as a context for implemented genetic circuits.

## 2 Results and discussion

### 2.1 Growth modulation and characterization by the engineered strain KT-TTX

*Pseudomonas putida*, as the rest of eubacteria, code for a single RNAP in the genome that is in charge of the full transcription activities of the cell. Each core enzyme is composed by two *α*, one *β* and one *β*’ subunits, coded by rpoA (PP 0479) gene and *rpoBC* operon (PP 0447-PP 0448), respectively [32]. Transcription of *rpoBC* operon is controlled by the activity of promoters pL11, which also controls *rplKAJL* ribosomal genes operon, and p1, placed next to attenuator PP mr07, which isolates rpoBC transcription from the ribosomal genes and is directly in charge of the regulation of transcription of *β* and *β*’ subunits [33] (Figure 1A). Additionally, *ω* subunit provides stability to the core complex, while specific *σ* subunits complete the holoenzyme complex. Driven by the *σ* subunit, the RNAP holoenzyme is capable of recognizing a selection of functionally related promoters and initiate transcription [34]. *α* subunit is produced constitutively and in excess, so the RNAP concentration is regulated by the amount of *β* and *β*’ subunits available in the cytoplasm [35] (Figure 1B). When higher levels of transcription are needed, *rpoBC* coded genes are transcribed in higher quantity, leading to a rise of the assembled *α*-*α*-*β*-*β*’, and therefore a higher availability of functional RNAP. In *E. coli*, pL11 and p1 respond to a variety of stimuli, like the stringent response triggered by the limitation of nutrients when cells are under a fast growth regime, orchestrated by the alarmone ppGpp [33, 36].

**Figure 1.**
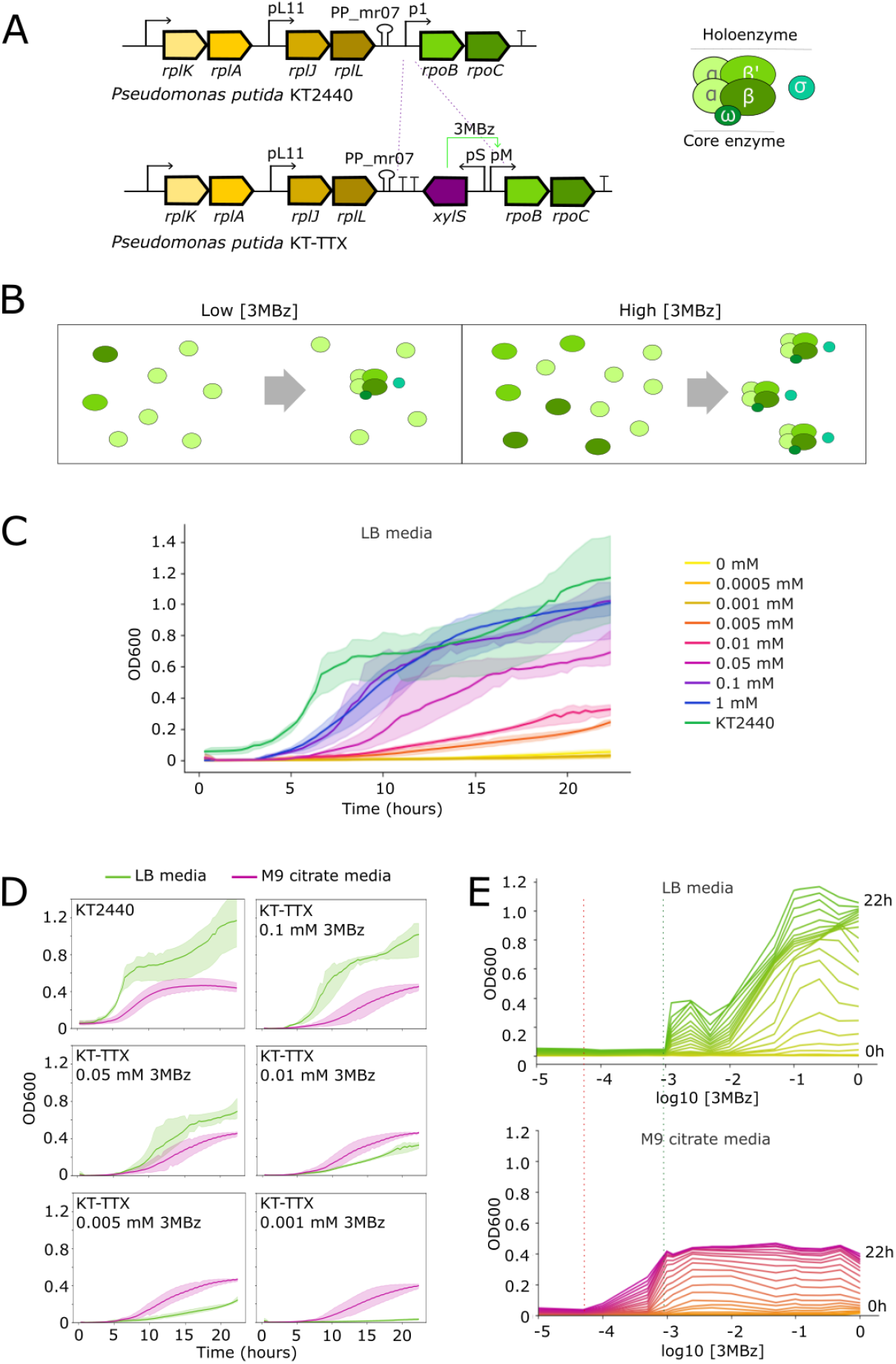
*Pseudomonas putida* KT-TTX produces RNAP under the control of XylS-Pm expression system. **A.** Genetic insertion of XylS transcription factor and its cognate promoter Pm next to *rpoBC* operon. This expression system is replacing native p1 promoter for the creation of the 3MBz growth responsive strain KT-TTX. *rpoB* and *rpoC* genes code for *β* and *β*’ subunits respectively that together with 2 *α* subunit and an *ω* subunit (core enzyme) conform the RNAP complex that is guided by specific *σ* subunits (holoenzyme) for the transcription of functionally related genes. **B**. In *P. putida* KT-TTX, low 3MBz levels keep scarce production of *β* and *β*’ subunits, restricting the assembly of RNAP to low concentration, and therefore keeping to the minimum general transcription activity of the strain. When 3MBz is increased, production of *β* and *β*’ subunits allow a higher RNAP assembly rates, having as a result increased transcription activity. **C**. Growth curves of KT-TTX in LB media under different 3MBz concentrations for 22 hours incubation. **D**. Growth curves comparison of *P. putida* KT2440 *wt* strain (first top left graph) and engineered KT-TTX (rest of graphs) for LB and M9 citrate media at specific 3MBz concentrations. In each graph it is noticeable how, at low levels of RNAP, M9 citrate cultures show an advantage over LB cultures, while at higher RNAP levels, LB media allows a faster and further growth of the strain. **E**. Growth response of KT-TTX to 3MBz at all tested times in LB and M9 citrate media. Red dashed line marks the growth turning point in M9 media, while green dashed line indicates this same growth change in LB media.

In order to control RNAP production at will, we have replaced p1 of *rpoBC* promoter by the expression system XylS-Pm (Figure 1A) and isolated rpoBC from pL11 regulation by adding a couple of transcriptional terminators (rpoC and T1) in tandem with the PP mr07 attenuator. Therefore, we managed to put transcription of *β* and *β*’ subunit under the control of chemical inducer 3-methyl benzoate (3MBz), used in *Pseudomonas* for tight control of gene expression, following a similar approach of previous works in *E. coli* [20], creating *Pseudomonas putida* KT-TTX strain. Different concentrations of 3MBz were tested in order to describe the dose response, dynamic range and operability of the system under two types of media: LB as an example of rich media and M9 citrate as a defined minimum media. Increasing concentrations of the inducer 3MBz generated a gradual response of growth observed by means of cell density values (measured by classical OD600) and growth rate for both LB and M9 citrate media (Figure 1C and S1). Despite the induction effect is reflected in the density levels of both media, different sensitivities to the inducer concentration can be found for each carbon regime. While M9 citrate media cultures needed a concentration of 5 x 10^*−*4^ mM to trigger growth, a similar phenotype was not observed in LB until 1 x 10^*−*2^ mM of 3MBz was reached (Figure S1). In fact, despite *P. putida* KT2440 divides faster and reaches higher OD600 values in LB media compared to M9 citrate, *P. putida* KT-TTX growth phenotype showed advantage in M9 media compared to LB media up to a concentration of 1 x 10^*−*3^ mM of 3MBz. When the inducer concentration was increased to 5 x 10^*−*3^ mM, the growth profile of LB vs M9 media switched, and LB cultures overgrew those at M9 (Figure 1D). Furthermore, (Figure 1E) shows the different inducer dose response of the strain in M9 media compared to LB at all the measured times. 3MBz dose triggering growth was slightly higher than an order of magnitude difference for both media. This observation suggests that machinery needed for citrate consumption, the only carbon source present in this M9 media, might cost less to produce in terms of transcription in an environment where RNAP availability is restricted to low levels. This might be related to previous works in which it was determined that more copies of rRNA were needed in *E. coli* in order to grow on rich media than in minimal media [37]. On the contrary, LB is a richer media and needs a more complex metabolism for the consumption of all the carbon components present in the media. This translates to higher demands on RNAP, which has to transcribe a higher number of metabolic genes. This may lead to a later response to the inducer in terms of concentration. However, once the critical RNAP availability is reached, the richness of LB allows culture density levels that are beyond the levels that M9 citrate can maintain.

### 2.2 Growth control leads to modifications of noise and reproducibility of gene expression

Manipulation of the production of the only RNAP present in the bacterial cell is expected to have consequences not only on the growth rate of the bacteria but also in the expression of any implemented gene or circuit of interest. This impact may vary from the final concentration of the protein of interest to the noise expression profile due to the well known involvement of transcription into that parameter. It has previously been reported that transcription rate of a gene, when translation keeps constant, affects to this noise profile of gene expression [38]. In order to check how RNAP availability can affect to the expression of a heterologous protein, we transformed our KT-TTX and the *wt* strain KT2440 with the pSEVA2313G plasmid (Figure 2A), coding for the msfGFP reporter gene under the pem7 constitutive promoter. The transformation outcome generated the strains KT-TTX-GFP and KT2440-GFP, respectively (see Table 1). GFP production was characterized in terms of fluorescence brightness and noise of gene expression under same two different media, LB and M9 citrate, at 16 and 22 hours of cultivation. Selected times corresponded to a time point where the 3MBz-induced growth levels showed higher growth differences (early stationary phase, 16 hours) and when growth levels start to converge (late stationary phase, 22 hours), something that allows to check the evolution of those parameters with time and growth. Four selected 3MBz concentrations were tested, corresponding to those that allowed 4 growth differentiated states at LB according to previous results: 0.005 mM, 0.01 mM, 0.5 mM and 0.1 mM 3MBz. Cytometry analysis was used to check on the status of other parameters related to bacterial growth, as a comparative and qualitative observation of cell sizes using FSC-A values.

**Table 1:**
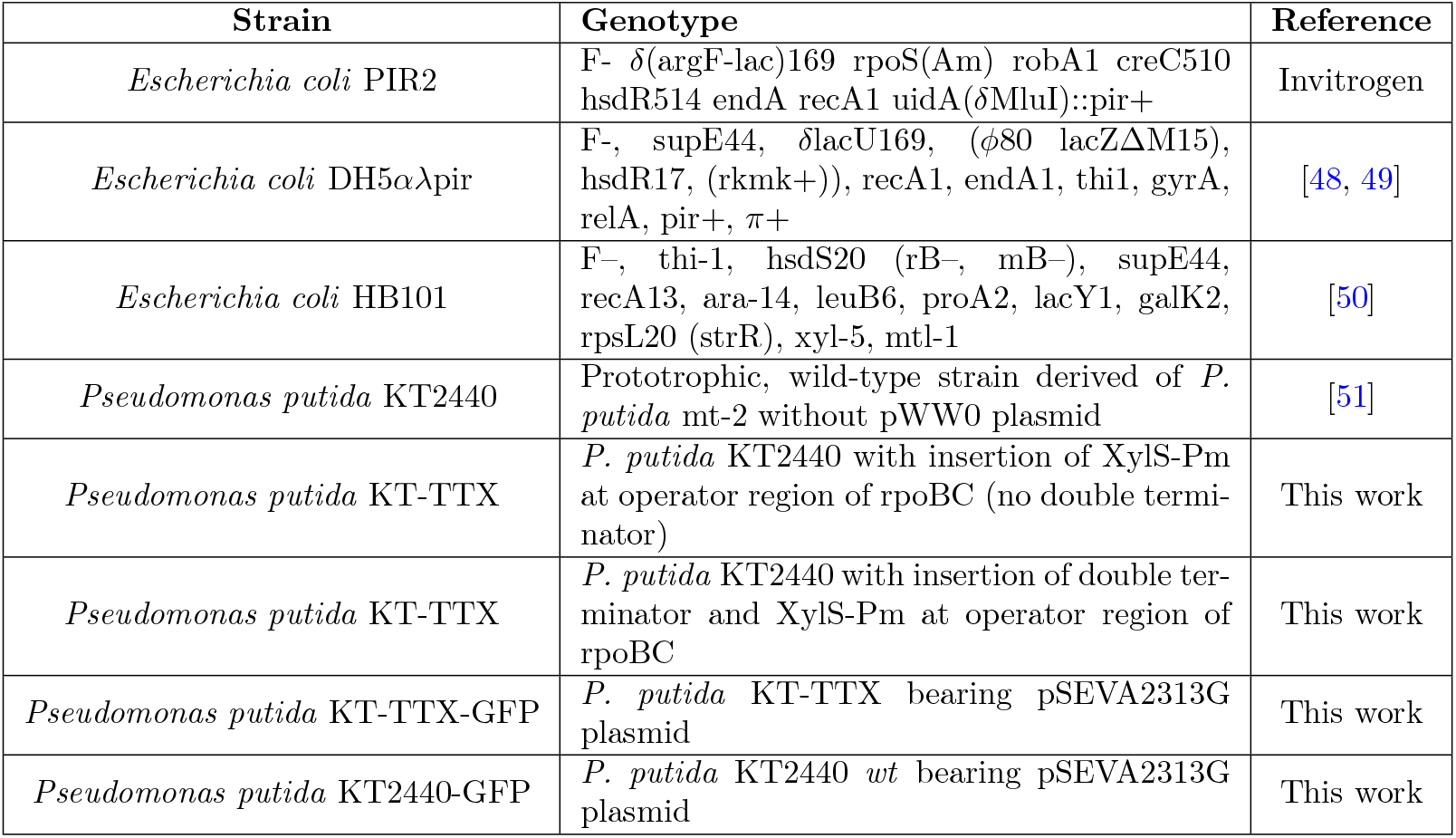
List of strains used in this work.

**Figure 2.**
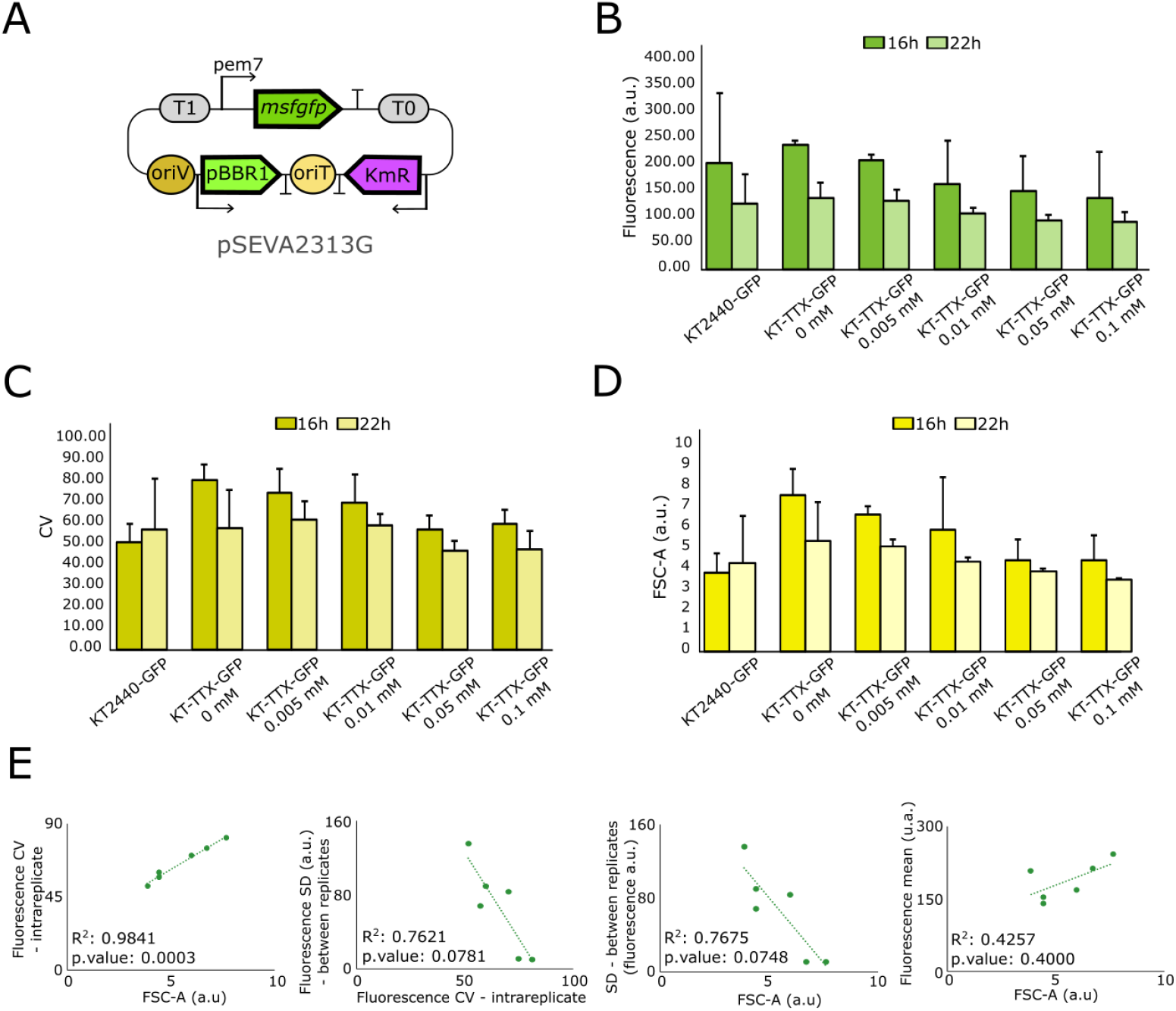
*Pseudomonas putida* KT-TTX coding for a constitutive expression of GFP allows deeper analysis of the influence of growth status over gene expression. Experiments shown in this picture were performed in LB, and three biological replicates per condition were plotted. **A**. Plasmid bearing msfGFP reporter gene under the constitutive control of pem7 promoter used to build KT-TTX-GFP and KT2440-GFP strains. **B**. Fluorescence reached by the *wt* KT2440-GFP and the engineered background KT-TTX-GFP at different concentrations of 3MBz at 16 and 22 hours. **C**. Noise of msfGFP expression (represented by the statistic CV) for the same conditions illustrates the impact of the RNAP availability over it. **D**. Indirect and qualitative report of cell sizes by FSC-A measurement under the same conditions show that lower amounts of 3MBz cause bigger cell sizes, despite cultures are less populated in terms of density. **E**. From left to right, correlations between noise and size, SD between replicates and intrareplicate noise, size and SD between replicates, and size and average fluorescence. This data suggests the interrelation of these three parameters modulated by the RNAP control through chemical induction.

Overall, RNAP induction demonstrated to have an impact over fluorescence expression levels, noise and (unrelated with the expression of a heterologous protein) bacterial size (Figure 2B-D and S2). Growth induction seemed to reduce GFP fluorescence per cell both in LB (Figure 2B) and M9 citrate media (Figure S2A), as nutrient-modulated growth model impacts on expression of a constitutive neutral gene [39]. This observation was linked to a reduction of individual cell size with higher concentrations of 3MBz (Figure 2D and S2C). As data suggests, induction of RNAP boosts cell division, populating cultures but reducing the average size of cells. Under low concentrations of 3MBz, bacteria kept slightly swollen, hosting more GFP molecules in their cytoplasm and rising fluorescence values, but without reaching the division point. These results were clearer at 16 hours incubation than at 22 hours, when culture densities started to be equal, so density levels and cell sizes appeared correlated and indirectly proportional. Therefore, we can hypothesize that at low levels of 3MBz and due to Pm leakiness, residual amounts of RNAP could be transcribing up to a level that keeps cells alive and slightly growing, but under the limit point in which division is triggered. When 3MBz increase RNAP concentration, growth and division cycle approach to *wt* levels and bacteria restore their phenotype in terms of density, division, size and average GFP expression.

It is important to remark that, with the decrease of size and fluorescence mean, noise of gene expression depicted by Coefficient of Variation (CV) also descends (Figure 2C and S2B). This is consistent with previous observations that assign higher noise levels to lower division rates [40]. Also, general noise levels in LB are lower than in M9 citrate, corresponding to poorer nutrient media and the possible need for cells of diversification strategies in those conditions. In previous experiments performed in *E. coli*, it was observed that RNAP concentration was also responsible of extrinsic noise but not of intrinsic noise [27], so although it was not directly measured, we infer that through RNAP manipulation extrinsic noise levels of KT-TTX heterologous gene expression may be changing. Furthermore, CV calculated values come from the noise in gene expression of one experimental replicate, and therefore one population. CV values are expected to be linked with Standard Deviation (SD) values of that same replicate (CV = SD/mean). However, it was observed that, with the increasing induction of growth, CV values of each replicate were decreasing while SD of fluorescence averages (calculated for the three replicates) were rising. Considering that reproducibility between replicates was decreasing with growth, we subjected the cytometry data of each replicate to a Kolmogorov-Smirnov statistical test to compare distributions and we found that replicates were more similar in terms of fluorescence distribution at lower RNAP induction (Supplementary Table S2). That means that noisier populations behave in a more predictable way in terms of reproducibility for the fluorescence mean, leading to lower SD values and more similar distributions between replicates. Induction of RNAP and lower levels of noise seem to impact on this reproducibility by varying mean fluorescence in every replicate but also cell sizes. This observation led to a further examination and, as Figure 2E shows, different correlations between cell size, noise intrareplicate and SD between replicates for our strain cultured in LB during 16 hours. Significance is stronger in the correlations of noise intrareplicate compared to size and SD between replicates, what suggests that the strain KT-TTX can be used to manipulate reproducibility and noise patterns at specific times and growth points. Commented correlations loose strength at 22 hours, again suggesting that differentiated growth states, induced by the expression of the RNAP, are influencing on fluorescence, noise and sizes. Figure S3 shows that, despite not all the correlations are significant, they are weaker in M9 citrate and at further times, when cultures have more similar growth rates. M9 media less significant results are probably due to the different sensitivity of KT-TTX culture to 3MBz in that media.

### 2.3 RNA polymerase availability shapes NOT gate circuit behavior under different dependence patterns

Growth control of KT-TTX through RNAP induction has shown a clear influence over the expression of a constitutive reporter and its different parameters. To further investigate the behavior of more complex circuits, we developed a mathematical model to predict the response of NOT gates under different growth conditions (see Materials and Methods).

Using data from the KT-TTX-GFP experiment, we estimated the RNAP availability as a function of 3MBz concentration (Figure S5A). A sensitivity analysis of this curve (Figure S5B) revealed three additional plausible RNAP vs 3MBz relationships (Figure S6). Using these four curves, the model predicted two distinct behaviors for the YFP vs IPTG response. The first predicted behavior corresponds to a non-inverting response, where YFP expression remains relatively flat across IPTG concentrations, with only the overall expression level shifting depending on 3MBz concentration. Three of the four RNAP vs 3MBz curves produced this non-inverting behavior, with only differences between low and high 3MBz depending on the specific RNAP curve considered (Figures S6A–C). The second predicted behavior corresponds to weakly inducer-dependent inverters, where the YFP vs IPTG curves display the expected inverting response (high YFP at low IPTG and vice versa), but with minimal differences between low and high 3MBz conditions (Figure S6D).

To experimentally test these predictions, KT-TTX strain was transformed with a collection of NOT gates that allowed to analyze the impact of growth manipulation over genetic circuits. These NOT gates are composed by the IPTG-responsive LacI-pTac system plus a set of TetR-homologous repressors and their cognate promoter supplying the inverting behaviour. The basic circuit structure controlling the expression of the reporter gene *yfp* appears in Figure S4. The collection of NOT gates was previously tested in the *P. putida wt* KT2240 host [6, 41], and the list of plasmids bearing the NOT gates used in this study can be consulted at Table 2. Growth induction levels in LB media were selected at concentrations of 0.005 mM (low), 0.05 mM (intermediate) and 0.1 mM (maximum growth) 3MBz. Simultaneously, all NOT gates were tested in a range of concentrations of the inducer IPTG from 0 (uninduced NOT gate expressing fluorescence) to 2 mM (totally induced NOT gate leading to *yfp* repression). Fluorescence expression was measured at 22 hours as NOT behavior is more distinguishable at late stationary phase and results could be contrasted with those obtained from constitutive GFP expression analysis performed in previous cytometry experiments.

**Table 2:** List of plasmids used in this work.

| Plasmid | Description | Reference |
| --- | --- | --- |
| pEMG | KmR, oriR6K, suicide plasmid with two I-SceI sites flanking the lacZα polylinker | [49] |
| pSWI | ApR, oriRK2, *xylS*, bearing a Pm → I-SceI transcriptional fusion | [55] |
| pRK600 | tra+, mob+, CmR, oriV ColE1, helper plasmid transformed into *E. coli* HB101 to assist conjugation between donor and receiver strain. | [56] |
| pTwist-XylSrpoBC | pTwist derivative, KmR, ori pUC, genetic fusion for fragment insertion of XylS-Pm at operator of *rpoBC* operon | Twist order |
| pEMG-XylSrpoBC | pEMG derivative, KmR, oriR6K, genetic fusion for fragment insertion of XylS-Pm at operator of *rpoB-rpoC* operon | This work |
| pEMG-XylSrpoBCTT | pEMG-XylSrpoBC derivative, double terminator inserted before XylS-Pm for a better isolation | This work |
| pSEVA2313G | KmR, ori pBBR1, pem7 → msfGFP | [57] |
| pS221-QACR-Q1 | pSEVA221 derivative, KmR, oriRK2, NOT gate QACR-Q1, *yfp* | [6] |
| pS221-QACR-Q2 | pSEVA221 derivative, KmR, oriRK2, NOT gate QACR-Q2, *yfp* | [6] |
| pS221-Srpr-S1 | pSEVA221 derivative, KmR, oriRK2, NOT gate Srpr-S1, *yfp* | [6] |
| pS221-Srpr-S2 | pSEVA221 derivative, KmR, oriRK2, NOT gate Srpr-S2, *yfp* | [6] |
| pS221-Srpr-S3 | pSEVA221 derivative, KmR, oriRK2, NOT gate Srpr-S3, *yfp* | [6] |
| pS221-Srpr-S4 | pSEVA221 derivative, KmR, oriRK2, NOT gate Srpr-S4, *yfp* | [6] |
| pS221-AmeR-F1 | pSEVA221 derivative, KmR, oriRK2, NOT gate AmeR-F1, *yfp* | [6] |
| p221HlyllR-H1 | pSEVA221 derivative, KmR, oriRK2, NOT gate HlyllR-H1, *yfp* | [6] |
| p221LitR-L1 | pSEVA221 derivative, KmR, oriRK2, NOT gate LitR-L1, *yfp* | [6] |
| p221LmrA-N1 | pSEVA221 derivative, KmR, oriRK2, NOT gate LmrA-N1, *yfp* | [6] |
| p221PhIF-P2 | pSEVA221 derivative, KmR, oriRK2, NOT gate PhIF-P2, *yfp* | [6] |
| pS221PsrA-R1 | pSEVA221 derivative, KmR, oriRK2, NOT gate PsrA-R1, *yfp* | [6] |
| pS221BetI-1 | pSEVA221 derivative, KmR, oriRK2, NOT gate BetI-1, *yfp* | [6] |
| pS221BM3R1-B1 | pSEVA221 derivative, KmR, oriRK2, NOT gate BM3R1-B1, *yfp* | [6] |
| pS221BM3R1-B2 | pSEVA221 derivative, KmR, oriRK2, NOT gate BM3R1-B2, *yfp* | [6] |
| pS221AmtR-A1 | pSEVA221 derivative, KmR, oriRK2, NOT gate AmtR-A1, *yfp* | [6] |
| pS221LCara-I1 | pSEVA221 derivative, KmR, oriRK2, NOT gate LCara-I1, *yfp* | [6] |

Most of the NOT gates could be categorized into three different cases depending on their performance across the different 3MBz concentrations. The first category comprises gates that do not exhibit NOT behavior at any 3MBz concentration (SrpR-S2, BM3R1-B1, BM3R1-B2, PhIF-P2, SrpR-S3, SrpR-S4 and BetI-E1) (Figures 3B, S6B and S6C), consistent with the non-inverting response predicted by the model (Figure 3A). Curiously, none of the tested clones behaved according to the original RNAP vs 3MBz curve derived from the KT-TTX-GFP experiment (Figure S6A). The second category includes gates that display NOT behavior at all 3MBz concentrations (LCara-I1, QacR-Q1 and QacR-Q2) (Figures 3D and S6D), in agreement with the weakly inducer-dependent inverters predicted by the model (Figure 3C). The third category consists of gates that only exhibit NOT behavior at high 3MBz concentrations but not at the lowest (HlyIIR-H1, LitR-L1, AmtR-A1) (Figures 3F and S7A), a group we refer to as strongly inducer-dependent inverters. This third behavior was not predicted by the model. To simulate such a response (Figure 3E), we had to assume different RNAP vs 3MBz relationships for each promoter controlling LacI, R and YFP expression (Figure 3E, left). Specifically, the curve for the LacI promoter corresponds to the non-inverting behavior (Figure 3A, left), whereas the curves for the R and YFP promoters correspond to the weakly inducer-dependent inverters (Figure 3C, left). Two additional clones (LmrA-N1 and SrpR-S1) were included in this third category (Figure S7A), despite showing NOT behavior at low 3MBz. This classification is justified because their YFP vs IPTG curves can be explained by the same RNAP vs 3MBz relationships as HlyIIR-H1, LitR-L1 and AmtR-A1. Following the same strategy of assuming promoter-specific RNAP vs 3MBz relationships, we were also able to explain the behavior of an additional clone, PsrA-R1 (Figure S7B), which did not fit into any of the three categories described above. However, the behavior of clone AmeR-F1 (Figure S7C) could not be reproduced by the model. In addition, gates corresponding to the first and third case (Figures 3B and F) displayed higher YFP expression levels under low 3MBz induction. These higher levels of fluorescence at lower growth rates align with those results obtained with the constitutive expression of GFP and therefore could be partially due to increased size and fluorescence at those conditions.

**Figure 3.**
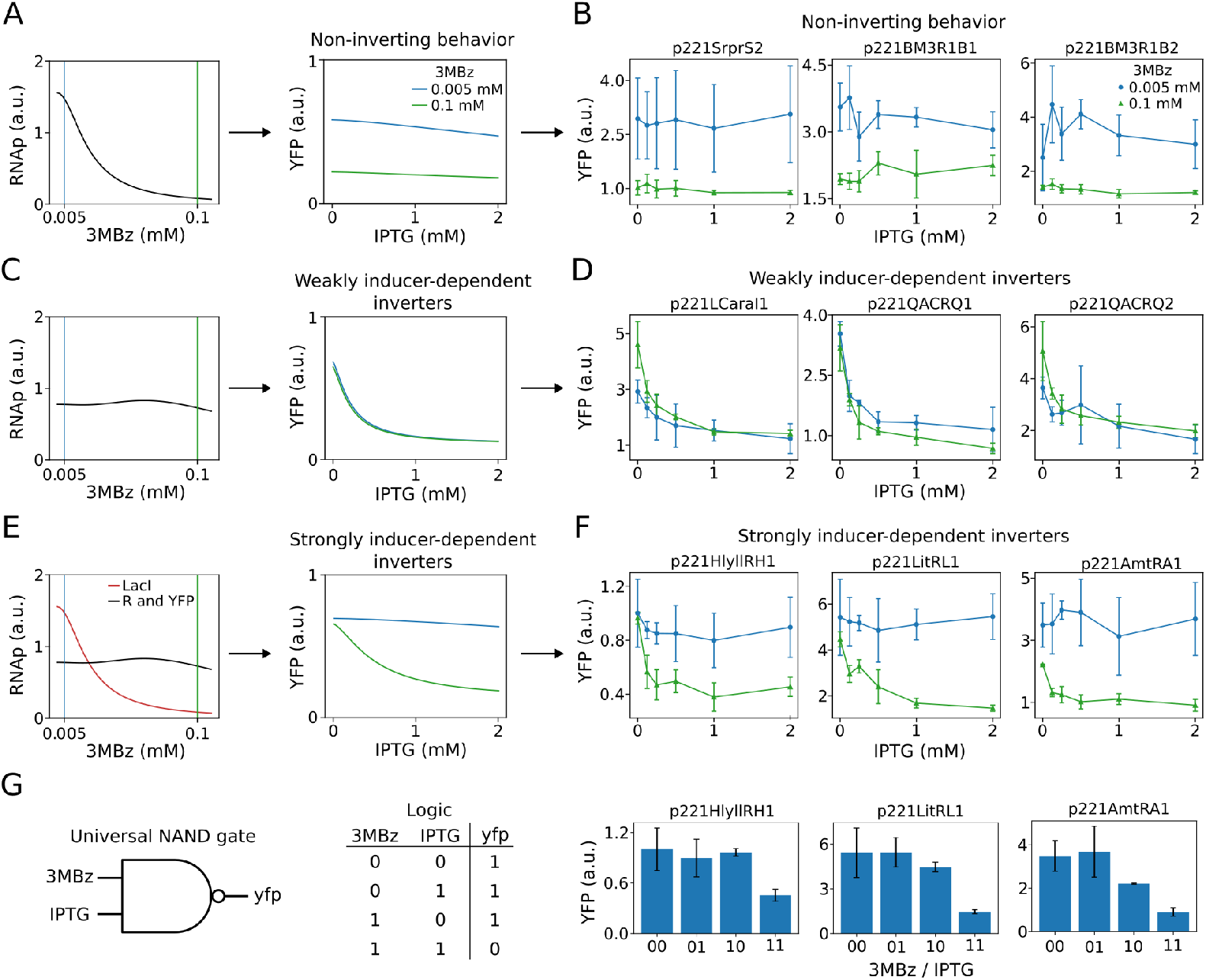
NOT gate behavior under growth-controlled conditions: model predictions and experimental validation. **A.** Model prediction for the non-inverting response. Left: assumed RNAP vs 3MBz relationship. Right: predicted YFP vs IPTG curves at low (0.005 mM) and high (0.1 mM) 3MBz concentrations. **B**. Experimental data for NOT gates displaying non-inverting behavior (see Figure S6C for additional clones). **C**. Model prediction for weakly inducer-dependent inverters. Left: assumed RNAP vs 3MBz relationship. Right: predicted YFP vs IPTG curves. **D**. Experimental data for NOT gates displaying weakly inducer-dependent inverting behavior. **E**. Model prediction for strongly inducer-dependent inverters. Left: promoter-specific RNAP vs 3MBz relationships assumed for LacI, R and YFP promoters. Right: predicted YFP vs IPTG curves. **F**. Experimental data for NOT gates displaying strongly inducer-dependent inverting behavior (see Figure S7A for additional clones). For simplicity, only the lowest (0.005 mM) and highest (0.1 mM) 3MBz concentrations are shown. Complete data including the intermediate concentration (0.05 mM) are presented in Figures S6 and S7. **G**. Emergent NAND gate behavior from panel F when 3MBz and IPTG are considered jointly as inputs. Left: schematic NAND gate with 3MBz and IPTG as inputs and YFP as output. Middle: corresponding truth table. Right: bar plots showing YFP levels for HlyIIR-H1, LitR-L1 and AmtR-A1 clones. The x-axis represents the four input combinations (3MBz/IPTG), where the first bit corresponds to 3MBz (0 = 0.005 mM, 1 = 0.1 mM) and the second bit to IPTG (0 = 0 mM, 1 = 2 mM). High YFP output is observed for all combinations except when both inputs are high (11).

Overall, these results show that the interplay between growth control and genetic circuit behavior can be captured by mathematical modeling through the consideration of promoter-specific RNAP availability. However, the inability of the model to reproduce the behavior of certain clones, such as AmeR-F1, suggests a more complex host-circuit interaction that require further investigation.

In addition to characterizing the growth-dependent behavior of individual NOT gates, the strongly inducer-dependent inverters revealed an emergent logic operation when 3MBz and IPTG were considered jointly as inputs. Specifically, the clones HlyIIR-H1, LitR-L1 and AmtR-A1 displayed high YFP expression for all input combinations except when both inducers were present at their highest levels, resulting in a low-output state. This pattern corresponds to a NAND logic function, as illustrated in Figure 3G. The emergence of a NAND gate is particularly relevant because NAND is a *universal* logic gate: any Boolean function can in principle be constructed by appropriate combinations of NAND operations. Moreover, using the growth control signal as input for a logic operation implies that the physical body of the cell deals with part of the information processing of the function. In other words, this is an example of morphological computing, an unconventional computing paradigm that deserves further attention, since living matter have features not found in classical hardware. Thus, the fact that growth-dependent modulation of RNAP availability naturally gives rise to a NAND-like behavior underscores not only the versatility of the host–circuit interaction but also its potential for enabling more sophisticated computation in engineered cells under growth-controlled conditions.

## 3 Discussion

Genetic circuits do not run in isolation. Rather, they are tightly connected to the cellular context in many ways, and their performance is modulated by this connection. Here, we engineered a tool for shifting the cellular context into one of many stable configurations, thus changing the nature of these connections (the so-called contextual dependencies). To this end, we implemented growth control on the soil bacterium *Pseudomonas putida* KT2440—a well-known synthetic biology workhorse [42]—by controlling RNA polymerase (RNAP) levels. This impacts both the growth rate and the internal workings of this bacterium simultaneously [43, 44].

Controlling bacterial growth is a key objective in the engineering of biology, but its many unknown effects can lead to the suboptimal use of any tool enabling this type of manipulation. Among the possible options to control this parameter, external induction of RNAP production was previously tested in *E. coli* [20] with a LacI expression system. That system, based on the de-repression of LacI by IPTG, required several copies of the *lacI* gene spread through the genome to circumvent natural genetic disruption.

The proposed *P. putida* KT-TTX strain, however, utilises an induction mechanism based on the activator XylS. This approach avoids the genetic disruption triggered by natural evolution that often complicates engineering efforts. The XylS-Pm expression system, controlling the native RNAP, allows the induction of different growth rates that are valuable for various purposes. For example, bringing growth rates to desired levels can be used for applications such as the design of artificial consortia, where different strain compositions need to be rebalanced according to functional requirements [13, 45], or for further exploration into uncoupling growth from production. Furthermore, the KT-TTX strain’s utility as a tool enables the deep study of cellular and gene expression parameters that cannot be explored at natural, self-regulated growth rates. The possibility of controlling growth at specific rates sheds light on the influence of RNAP concentration and consequent bacterial cell sizes on gene expression, noise, and reproducibility, revealing complex interplays. While we have discussed all these factors, it is not yet possible to weigh their individual impact at different growth speeds. However, it is indeed possible to assign differential context states that correlate to genetic circuit function.

Using a mathematical model, we were able to correlate growth to circuit function. We achieved this by characterising a library of NOT logic gates at several growth states. While it is true that growth rates did not homogeneously tune all logic gates in the same way (or rather, their contextual dependencies), we observed that this tuning is not as heterogeneous as initially expected. Indeed, we classified three tuning patterns within our library of gates. In other words, logic gates changed their performance in one of three distinct modes. This suggests that growth control, and in turn, control over circuit-host interplay, can be used as a reliable engineering tool for tuning circuit performance.

Finally, we highlighted those conditions under which a subset of circuits exhibited a NAND-like response, considering growth control as an input. This suggests the potential for growth modulation to enable not only universal logic, but also morphological computing in living cells. Although Boolean logic and other conventional approaches are still mainstream in genetic circuitry [46], the search for unconventional computing paradigms is becoming increasingly active [47]. Altogether, KT-TTX represents a significant opportunity for both basic research and derived applications in biological engineering and biocomputation.

## 4 Materials and methods

### 4.1 Culture media, conditions and strains

Strains used in this work are listed in Table 1. Both *Escherichia coli* or *Pseudomonas putida* strains were usually grown in Lysogeny Broth (LB) media. Additionally, *P. putida* was grown in M9 minimal media supplemented with citrate 0.2% (v/w) when this media was required. Antibiotics were added when needed at the following concentrations: kanamycin (Km) 50 *µ*g/mL, gentamicin (Gm) 10 *µ*g/mL, and ampicillin (Ap) 500 *µ*g/mL (concentration required for *P. putida*). Inducers were included at the indicated concentrations using serial dilutions stocks: 3-Methylbenzoate (3MBz; also known as m-Toluate or m-Toluic acid) at 500 mM, 50 mM, 5 mM and 0.5 mM. Isopropyl *β*-d-1-thiogalactopyranoside (IPTG) was used from a stock at 1mM in order to induce the NOT gates at 0.125, 0.25, 0.5, 1 and 2 mM.

### 4.2 DNA assemblies

For the preparation of DNA constructs, PCR was performed using Phusion (Thermo Fisher Scientific, Waltham, Massachusetts, USA) or Q5 (New England Biolabs, Ipswich, Massachusetts, USA) proof-reading enzymes. Primers (a full list can be checked in Supplementary Material, Table S1) were ordered to IDT (Coralville, Iowa, USA). The presence of DNA insertions in the genome was checked by PCR using PhirePlant enzyme (Thermo Fisher Scientific, Waltham, Massachusetts, USA). When PCR’s were cleaned for later constructs, Monarch PCR and DNA Cleanup kit (New England Biolabs, Ipswich, Massachusetts, USA) was used. Fragments of DNA were assembled either by isothermal assembly [52] (Gibson Assembly MasterMix, New England Biolabs, Ipswich, Massachusetts, USA) or classical digestion-ligation protocols [53] (restriction enzymes, T4 DNA Ligase and Quick Ligation kit from New England Biolabs, Ipswich, Massachusetts, USA). pEMGXylSrpoBC was built by digesting pTwist-XylSrpoB ordered plasmid (Twist Bioscience, California, USA) with enzymes SacI and BamHI, as well as vector pEMG used for chromosomal recombinations [49]. After proper ligation, the resulting plasmid pEMG-XylSrpoBC was used for the additional insertion of a double terminator by RF cloning technique [54] with primers *RF XylSrpoBTT F* and *RF XylSrpoBTT R* and a template sequence including an *rpoC* terminator ordered as a gBlock (named PP mr07-rpoC; Supplementary Table S2) to IDT (Coralville, Iowa, USA). Outcome plasmid pEMG-XylSrpoBCTT was used for delivery of the desired construct into the chromosome by three-parental conjugation using *P. putida* KT2440 as the recipient strain: XylS-Pm expression system was inserted in the regulatory region of the operon *rpoB-rpoC* (PP 0447-PP 0448), replacing its natural regulatory region. *E*.*coli* HB1010 bearing pRK600 plasmid was used as a helper strain to transfer plasmid from *E. coli* PIR2 carrying the pEMG-XylSrpoBCTT into *P. putida*. Candidate clones were used for amplification with *rplJ seq F* and *rpoB seq R* primers (see Table S1) and the PCR outcome was sent for sequencing to Macrogen (Seul, South Corea). Minipreps were performed using Monarch Plasmid Miniprep kit (New England Biolabs, Ipswich, Massachusetts, USA). Plasmids used in this study can be consulted in Table 2 2. The rest of replicative plasmids were transformed into *P. putida* by electroporation using 300 mM sucrose protocol [49].

### 4.3 Growth characterization

Cultures of *P. putida* KT2440 *wt* and *P. putida* KT-TTX were incubated in tubes with 5 mL of LB media at 30ºC with shaking for an O/N (16 h). *P. putida* KT-TTX precultures were always cultivated in the presence of 1mM of 3MBz in order to allow growth. After the O/N, cultures were washed to remove the inducer and they were resuspended in 5 mL of fresh LB without the inducer. 96-well plates with 200 *µ*L of LB or M9 citrate with the adequate antibiotics were inoculated with 0.5 *µ*L of the *P. putida* resuspension in clean LB. The set of 3MBz dilution stocks at 500 mM, 50 mM, 5 mM and 0.5 mM were used to generate the range of induction concentrations tested in this work. Density reads were performed using Varioskan LUX (Thermo Fisher Scientific, Waltham, Massachusetts, USA). For the growth curves, measurement conditions consisted in 1 read every 20 minutes during a total of 22 hours at an incubation temperature of 28ºC (to reduce evaporation of media) and intermittent shaking (5 seconds of medium shaking every 20 minutes).

### 4.4 Flow cytometry

LB precultures were set overnight and shaked at 30ºC. The following day, 5 *µ*L of preculture was used to inoculate 5 mL of LB or M9 citrate media with the corresponding concentration of 3MBz. After incubation at 30ºC with shaking for 16 or 22 hours, 100 *µ*L of cultures were pelleted and re-suspended in 1 mL of filtered PBS. Cultures were passed at a range of 800-1300 events per second in order to not to artifact noise measurements with higher speeds. Fluorescence, FSC-A and CV values were calculated for every replicate, and after 3 biological replicates, averages and SD values for the whole experiment were plotted.

### 4.5 NOT gates characterization

*P. putida* KT-TTX bearing the different NOT gates from reported list (see Table 2) were cultured in LB plus antibiotics for an O/N at 30ºC with shaking, including 1mM of 3MBz to allow growth as commented above. Following day, cultures were washed and resuspended in LB and 0.5 *µ*L were used to inoculate 200 *µ*L of LB or M9 citrate, dispensed both in 96-well plates with different concentrations of 3MBz (0.005, 0.05 and 0.1 mM) and IPTG (0, 0.125, 0.25, 0.5, 1 and 2 mM). Plates were placed at 30ºC with 400 rpm shaking for 22 hours. After the incubation time, a point measurement for OD600 and YFP fluorescence was taken using Varioskan LUX.

### 4.6 Mathematical Model

We developed a mathematical model to describe the dynamics of a three-gene regulatory circuit consisting of genes L, R, and Y. The circuit implements a cascade architecture where gene L is inhibited by IPTG, gene R is repressed by the active form of protein L, and gene Y is repressed by protein R. Additionally, the system’s transcription rates are modulated by the availability of RNA polymerase (RNAP), which depends on the concentration of 3MBz (*M*) in the medium.

Induction with 3MBz modulates the availability of the host RNAP for gene expression. We coarsegrain the effective RNAP concentration available for the circuit genes into a single variable *P* (*t*). Its temporal dynamics follow:

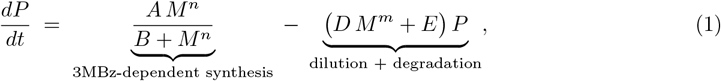

where the parameters capture distinct biological processes:

- *A, B*, and *n* parameterize the Hill-type induction response: *A* sets the maximal induction rate, *B* the half-saturation constant, and *n* the cooperativity;
- The effective loss rate *D M*^*m*^ + *E* increases with inducer concentration, where *D M*^*m*^ captures enhanced dilution due to faster growth at higher RNAP levels, and *E* represents basal dilution plus active degradation.

We assume a quasi-steady-state approximation in which the RNAP dynamics equilibrate faster than the circuit proteins. Setting 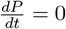 in Eq. (1) yields:

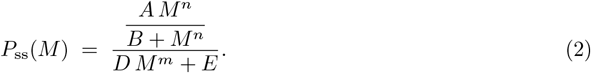

The parameter values were fitted based on the experiments performed with the KT-TTX-GFP strain (Figure S5A).

Different genes may compete differently for RNAP or have distinct promoter-RNAP affinities. We capture these effects through gene-specific parameter sets *p*_*i*_ = (*L*_*i*_, *A*_*i*_, *B*_*i*_, *n*_*i*_, *D*_*i*_, *m*_*i*_, *E*_*i*_) for *i ∈* {*L, R, Y*}. The normalized RNAP availability factors are:

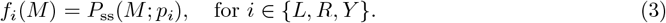

These dimensionless factors modulate the transcription rates in the circuit dynamics.

The temporal evolution of the NOT gate is described by the following system of ordinary differential equations (ODEs):

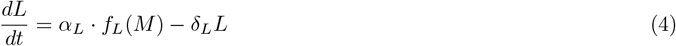

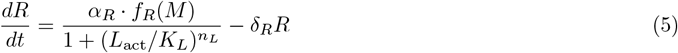

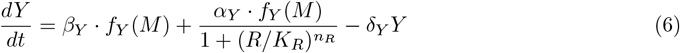

where:

- *L, R*, and *Y* represent the concentrations of the respective proteins
- *α*_*i*_ are the transcription rates for each gene
- *f*_*i*_(*M*) are the normalized RNAP availability functions
- *δ*_*i*_ are the protein degradation/dilution rates
- *β*_*Y*_ represents the basal transcription rate of gene Y
- The term 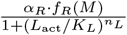 represents the production rate of protein R under repression by active L. The numerator (*α*_*R*_ *· f*_*R*_(*M*)) is the maximum transcription rate of R modulated by RNAP availability, while the denominator implements a Hill function modeling transcriptional repression: as active L concentration (*L*_act_) increases, the denominator grows and reduces R production. Parameters *K*_*L*_ and *n*_*L*_ represent the dissociation constant and Hill coefficient (cooperativity), respectively.
- The term 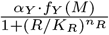 represents the inducible production rate of protein Y under repression by R. Similarly, the numerator (*α*_*Y*_ *· f*_*Y*_ (*M*)) is the maximum transcription rate of Y, and the denominator models repression by R through a Hill function. As R increases, Y expression decreases. Parameters *K*_*R*_ and *n*_*R*_ are the dissociation constant and cooperativity coefficient for this regulatory interaction.

The active form of protein L is modulated by IPTG according to:

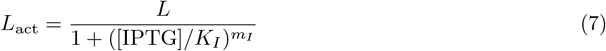

where *K*_*I*_ is the IPTG dissociation constant and *m*_*I*_ is the Hill coefficient for IPTG binding.

The parameter values for the NOT-gate model can be found in Table S4 of the Supporting Information.

## Acknowledgments

This work was supported by the the ECCO (ERC-2021-COG-101044360) Contract of the EU, and grants MULTI-SYSBIO (PID2020-117205GA-I00), and MULTI-SYNBIO (PID2023-152470NB-I00) funded by MICIU/AEI/ 10.13039/501100011033.

