## Supplementary Information for "Growth control as a central regulator for tuning the cellular context"

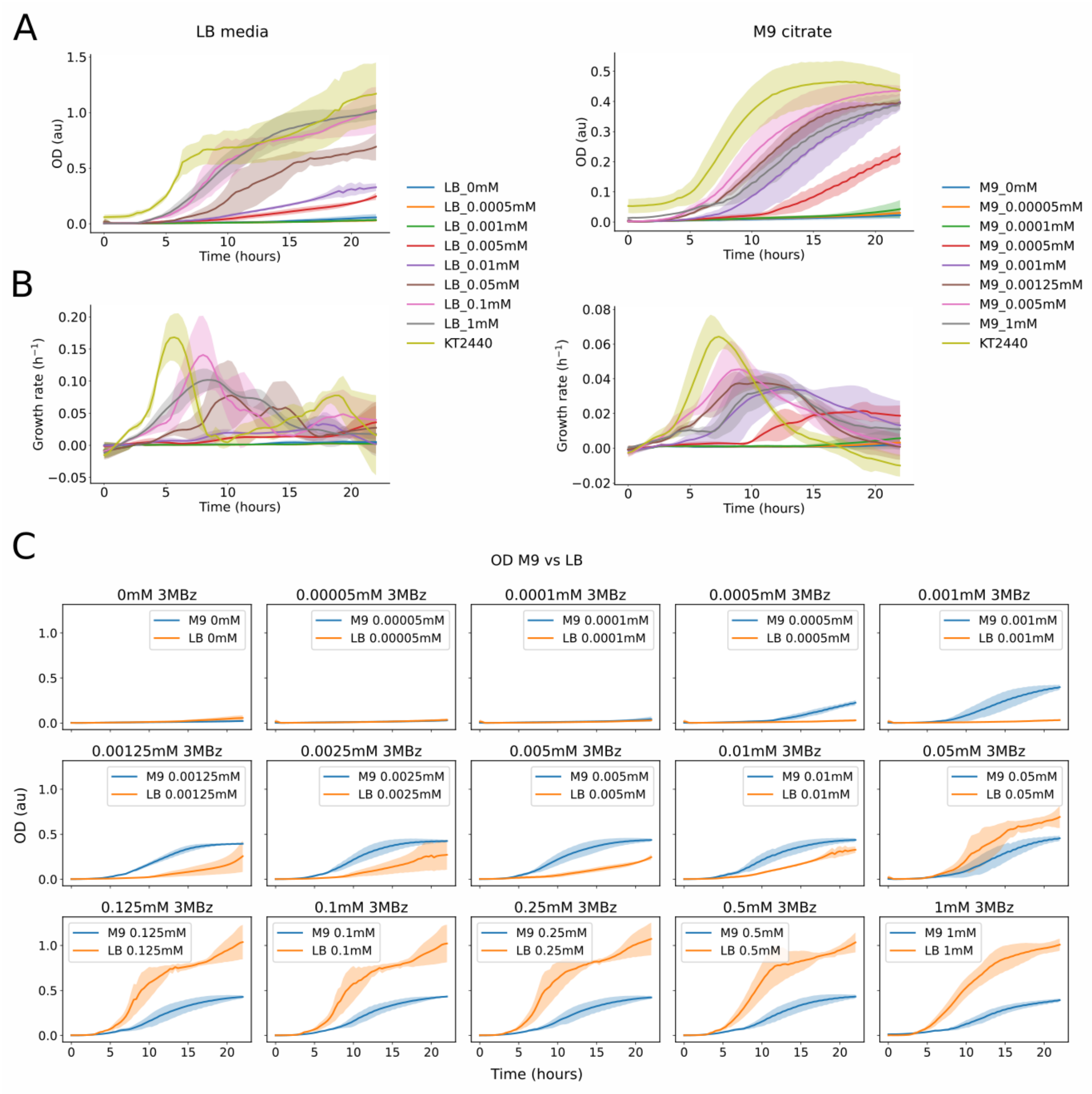


Figure S1: Figure S1: *Pseudomonas putida* KT-TTX growth response to the induction with 3MBz in LB and M9 media. **A.** Growth curves displaying cell density measured during 22 hours for both media at different 3MBz concentrations. **B.** Growth rates calculated for the same conditions. **C.** LB and M9 citrate differences in growth for each 3MBz concentration tested, depicted to evidence the differences between media at each inducer concentration.

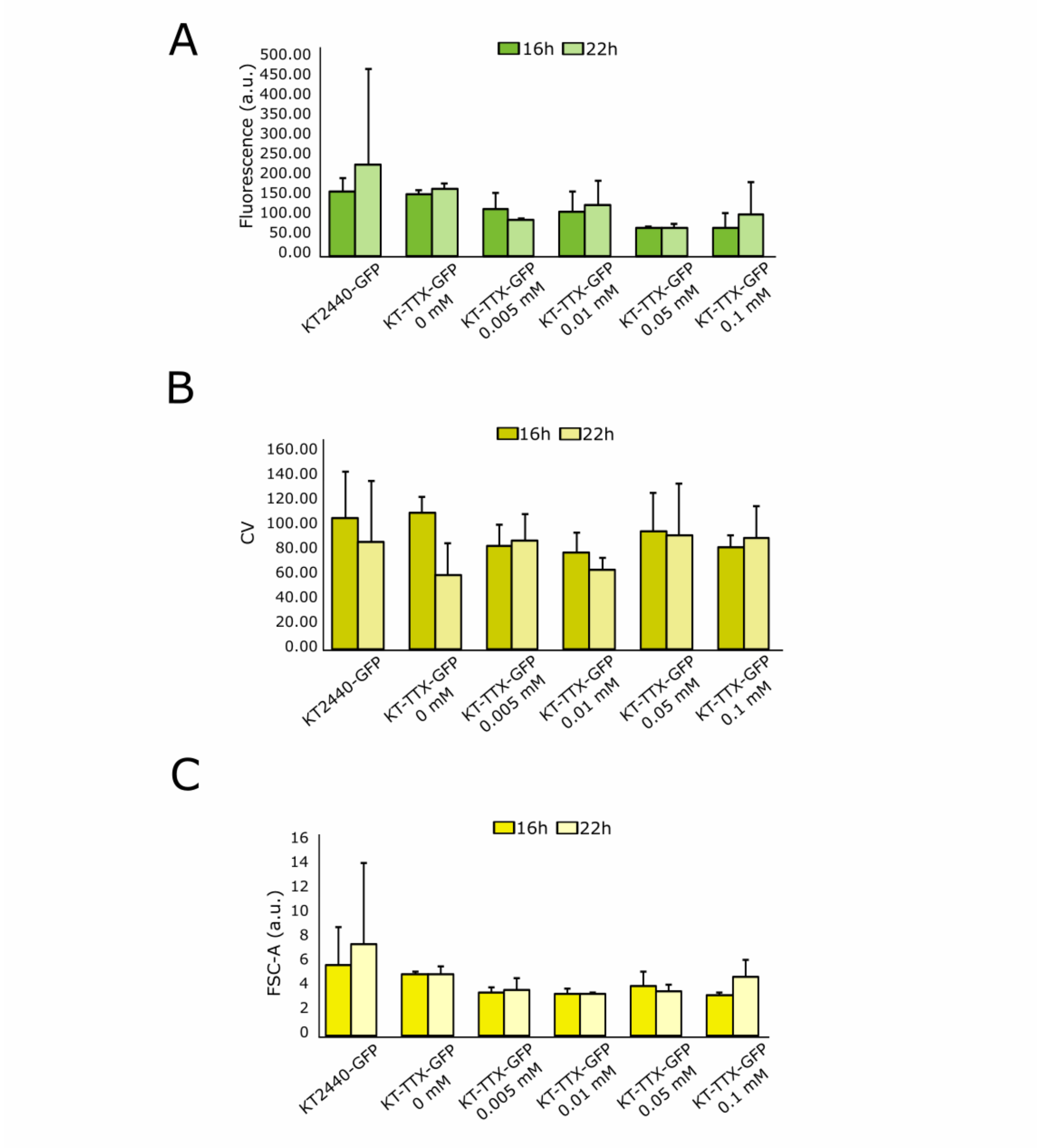


Figure S2: Figure S2: Analysis of the constitutive GFP fluorescence expression by KT-TTX-GFP at different growth induction levels and the *wt* control KT2440-GFP in M9 media at 16 and 22 hours. Three biological replicates were performed by condition. **A.** Average fluorescence measured by the cytometer for the strains and conditions tested. **B.** Average of the noise levels represented by CV values per replicate are plotted for the same conditions. **C.** Qualitative and indirect measurement by FSC-A values of cell sizes for the same experiments.

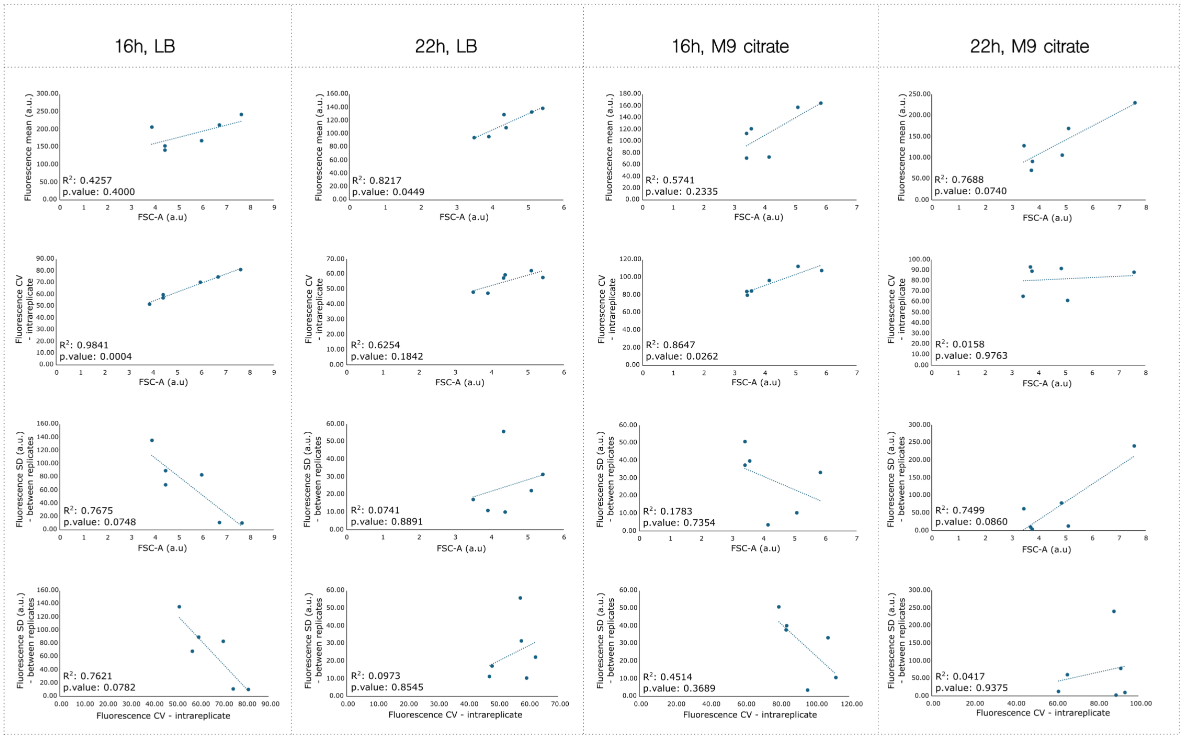


Figure S3: Figure S3: Correlations between size (FSC-A), noise (CV), reproducibility (SD) and fluorescence-mean for the conditions tested, for LB and M9 citrate media measured at 16 and 22 hours.

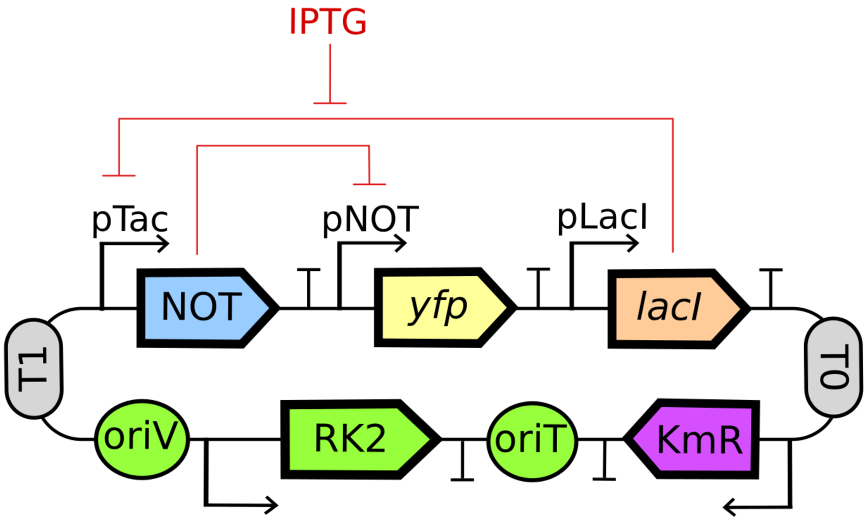


Figure S4: Figure S4: NOT gate circuit structure used in this study, beared by low copy number plasmid (pSEVA221 backbone) which were transformed into *P. putida* KT-TTX strain to study the interplay between induced growth status of the culture and circuit behaviour.

A

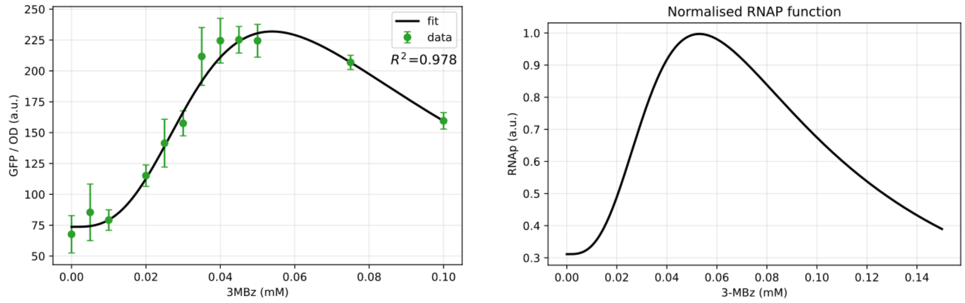


B

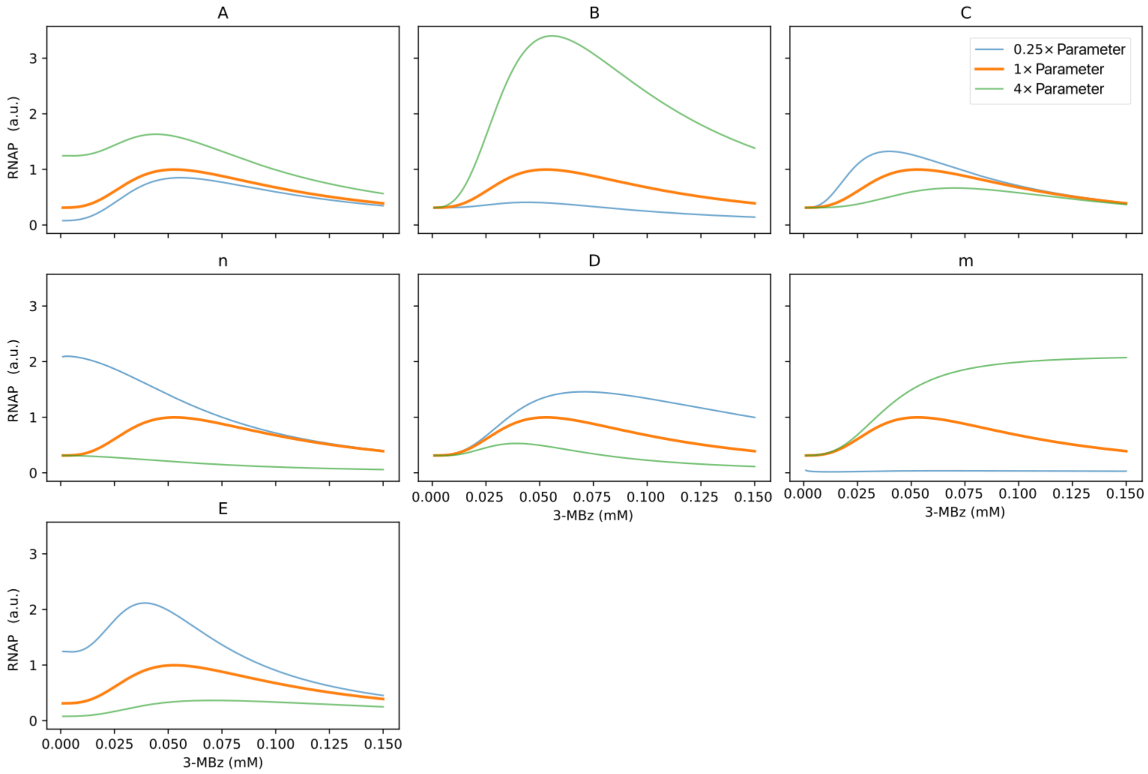


Figure S5: Figure S5: RNAP vs 3MBz curve fitting and sensitivity analysis. **A.** Left: fitting of the RNAP vs 3MBz curve to experimental data from the KT-TTX-GFP strain. Fitted parameters: $A = 104$, $B = 603$, $C = 1.0 \times 10^{-4}$, $n = 2.86$, $D = 251$, $m = 1.96$, $E = 1.41$. Right: normalized curve (maximum set to 1); normalization only affects $A$ and $B$, yielding $A = 0.439$ and $B = 2.54$. **B.** Sensitivity analysis of the normalized RNAP vs 3MBz curve. Each panel shows the effect of varying a single parameter ($A$, $B$, $C$, $n$, $D$, $m$ or $E$) by factors of 0.25 and 4 relative to the fitted value, while keeping all other parameters constant. This analysis reveals the characteristic RNAP vs 3MBz curve shapes that were later selected as representative scenarios for the simulations in Figure S6.

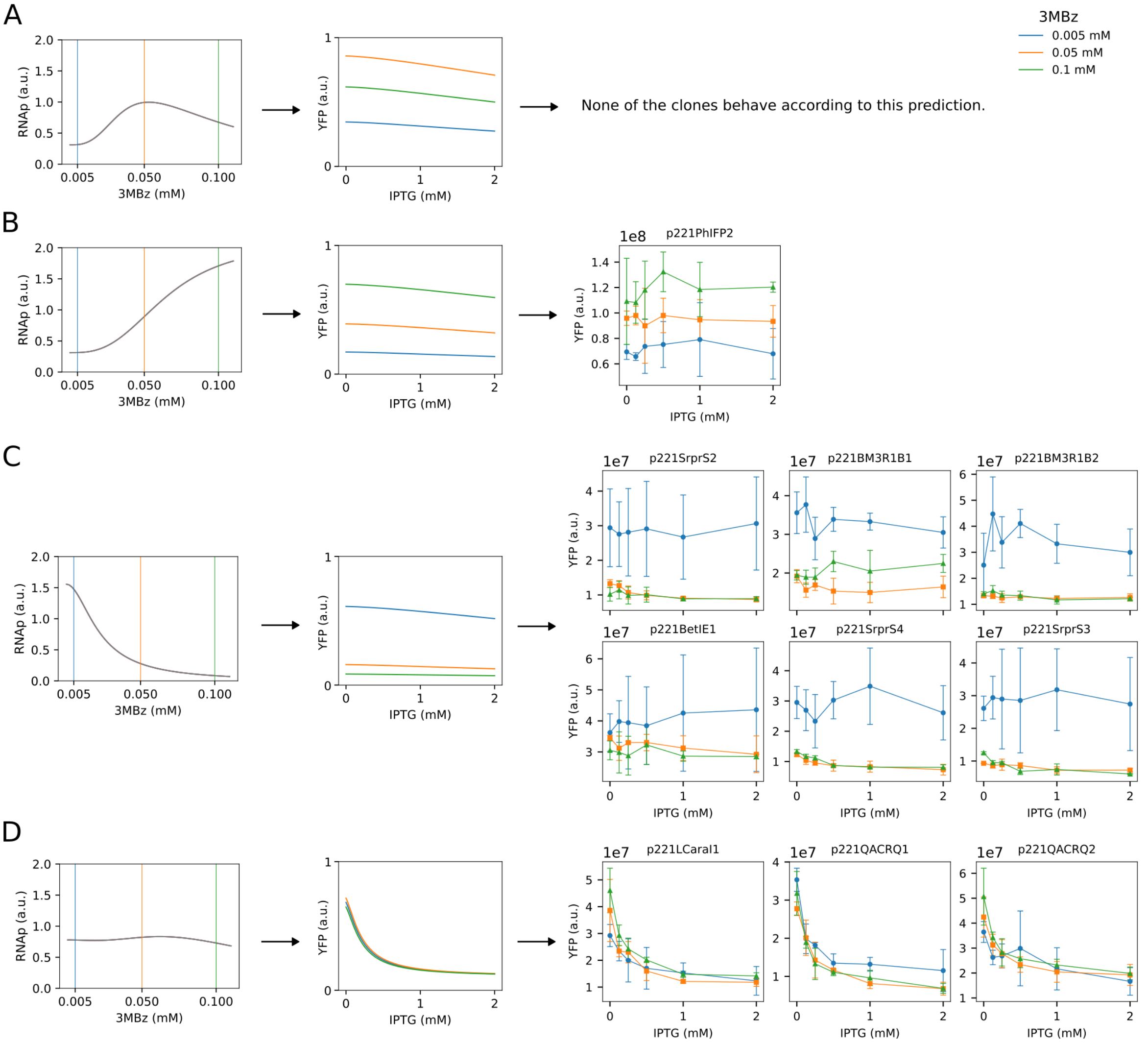


Figure S6: Figure S6: Complete NOT gate characterization: model predictions and experimental data for non-inverting and weakly inducer-dependent behaviors. **A.** Model prediction for the first non-inverting scenario (RNAP vs 3MBz curve derived from the KT-TTX-GFP experiment (Figure S5A)). Left: RNAP vs 3MBz curve. Center: predicted YFP vs IPTG curves. Right: None of the tested clones exhibited this predicted behavior. **B.** Model prediction for the second non-inverting scenario with an alternative RNAP vs 3MBz relationship (high 3MBz yields higher flat YFP curves). Left: RNAP vs 3MBz curve. Center: predicted YFP vs IPTG curves. Right: experimental data (1/17 clones matched this behavior). **C.** Model prediction for the third non-inverting scenario with an alternative RNAP vs 3MBz relationship (high 3MBz yields lower flat YFP curves). Left: RNAP vs 3MBz curve. Center: predicted YFP vs IPTG curves. Right: experimental data (6/17 clones matched this behavior). **D.** Model prediction for weakly inducer-dependent inverters. Left: RNAP vs 3MBz curve. Center: predicted YFP vs IPTG curves. Right: experimental data (3/17 clones matched this behavior). All panels display data for three 3MBz concentrations: 0.005 mM (low), 0.05 mM (intermediate) and 0.1 mM (high). The intermediate 3MBz concentration (0.05 mM) is included here because it provides an essential constraint for distinguishing among RNAP vs 3MBz curves that would otherwise be indistinguishable using only the low and high concentrations shown in Figure 3. Simulation parameters: the base RNAP vs 3MBz curve parameters $(A, B, C, n, D, m, E)$ are detailed in Figure S5; NOT gate parameters are listed in Table S4. For all scenarios, the same RNAP vs 3MBz parameters were assumed for all three promoters ($p_L = p_R = p_Y$). Panel-specific modifications: (A) original parameters, $K_R = 10^{-4}$ mM; (B) $(A, B, 4C, n, D, 3m, E)$, $K_R = 10^{-4}$ mM; (C) $(5A, 0.1B, C, n, 10D, m, E)$, $K_R = 10^{-4}$ mM; (D) $(5A, B, 5C, n, D, m, 2E)$, $K_R = 10^{-6}$ mM.

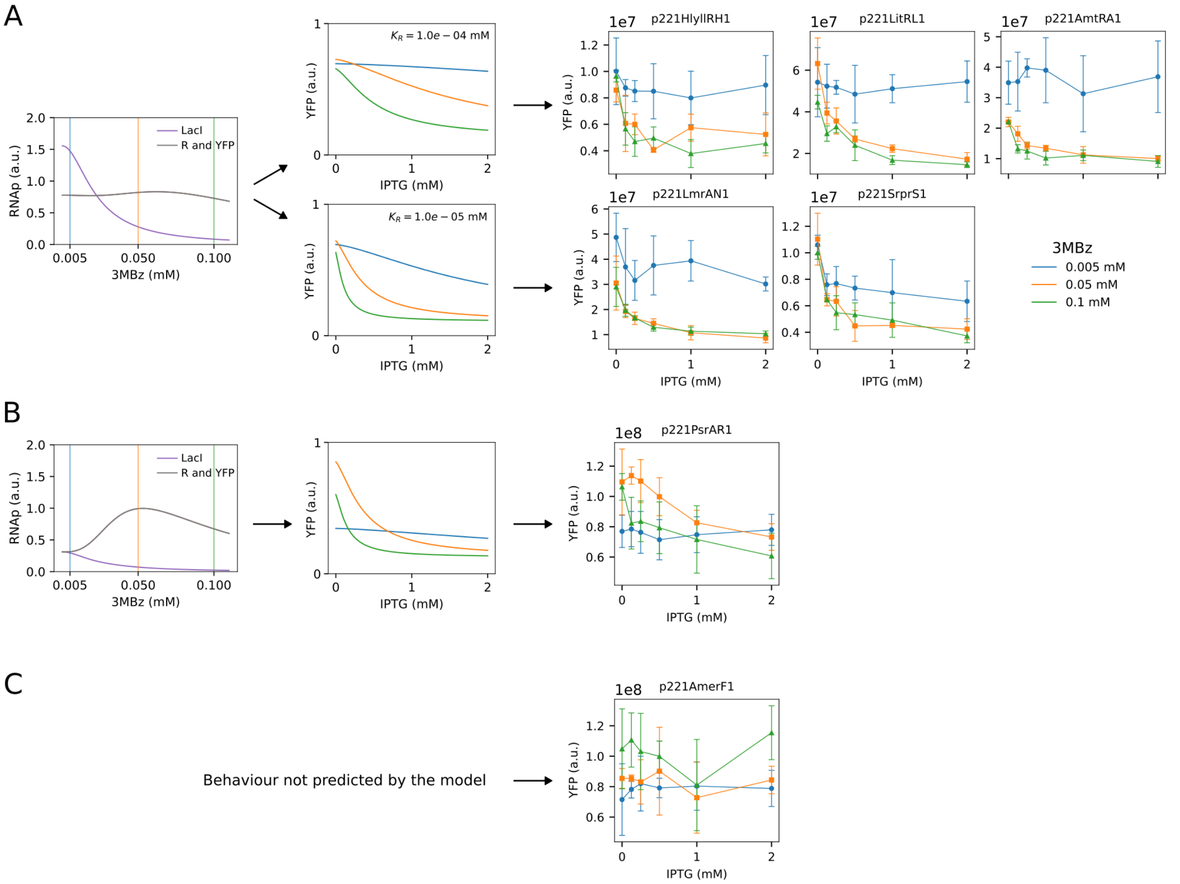


Figure S7: Figure S7: Complete NOT gate characterization: strongly inducer-dependent inverters and outlier clones. **A.** Model prediction for strongly inducer-dependent inverters. Left: RNAP vs 3MBz curves for LacI ($p_L$) and for R and YFP ($p_R = p_Y$). Center: predicted YFP vs IPTG curves. Right: experimental data (5/17 clones matched this behavior, including two clones—LmrA-N1 and SrpR-S1—that display NOT behavior at low 3MBz). **B.** Model prediction for an outlier clone (PsrA-R1) that did not fit into the three main categories but could be explained by assuming promoter-specific RNAP vs 3MBz relationships. Left: RNAP vs 3MBz curves for LacI ($p_L$) and for R and YFP ($p_R = p_Y$). Center: predicted YFP vs IPTG curves. Right: experimental data. **C.** Experimental data for clone AmeR-F1, whose behavior could not be reproduced by the model under any of the tested assumptions. All panels display data for three 3MBz concentrations: 0.005 mM (low), 0.05 mM (intermediate) and 0.1 mM (high). Simulation parameters: the base RNAP vs 3MBz curve parameters $(A, B, C, n, D, m, E)$ are detailed in Figure S5; NOT gate parameters are listed in Table S4. Panel-specific modifications: (A) $p_L = (5A, 0.1B, C, n, 10D, m, E)$, $p_R = p_Y = (5A, B, 5C, n, D, m, 2E)$, $K_R = 10^{-4}$ mM (except for LmrA-N1 and SrpR-S1, where $K_R = 10^{-5}$ mM); (B) $p_L = (A, 0.1B, C, n, 10D, m, E)$, $p_R = p_Y = (A, B, C, n, D, m, E)$, $K_R = 10^{-4}$ mM.

| **Primer** | **Sequence** | **Template** | **Final Construct / Usage** |
|---|---|---|---|
| RF XylSrpoB TT F | GGCGCTAAAGTCGAGCT GAAGTAATCTCGACCTT GCGATGCC | gBlock PP_mr07_rpoC | pEMG-XylSrpoBCTT |
| RF XylSrpoB TT R | GCCCTAGACAGCTGGGC GCGCCAGCGTGACAAAT GCTCTTTCC | gBlock PP_mr07_rpoC | pEMG-XylSrpoBCTT |
| rplJ seq F | TGGCTCGTACTCTGGCA GCC | *P. putida* genome | Amplification for sequencing |
| rpoB seq R | TGTCCATGACTGCAACT TCACC | *P. putida* genome | Amplification for sequencing |

Table S1: List of primers used in this study.

| gBlock | Sequence | Final Construct |
|---|---|---|
| PP_mr07_rpoC | tcTCGACCTTGCGATGCCAGCCTGAGCGTTGAGCG<br>AAAGGCTGATGGCTGGTGGCTTATGCCACCGGCC<br>TTTTTCCGTTATTGGTGGCCGGTCAGGCCGGTCC<br>CGATAACGAGCTGCAAGACACCCATGTGGGTGGC<br>GCAAACCAAGGGGTTTGCACGATTTTCTGGCTGC<br>TCCCGCCGGGAGAAGCCAAACAAGCAGGTGACCA<br>AGGTAATCGTTAATCCGCAAATAACGTAAAAACCC<br>GCTTCGGCGGGTTTTTTTATGGGGGGAGTTTAGG<br>GAAAGAGCATTTGTCAcgct | pEMG-<br>XylSrpoBCTT |

Table S2: gBlock ordered and used in this study.

| Strain and condition | Comparison | KS test value |
|---|---|---|
| KT-TTX-GFP 0.005 mM 3MBz | Replicate 1 vs replicate 2 | 0.1798 |
| KT-TTX-GFP 0.005 mM 3MBz | Replicate 1 vs replicate 3 | 0.2061 |
| KT-TTX-GFP 0.005 mM 3MBz | Replicate 2 vs replicate 3 | 0.1426 |
| KT-TTX-GFP 0.01 mM 3MBz | Replicate 1 vs replicate 2 | 0.5883 |
| KT-TTX-GFP 0.01 mM 3MBz | Replicate 1 vs replicate 3 | 0.4715 |
| KT-TTX-GFP 0.01 mM 3MBz | Replicate 2 vs replicate 3 | 0.1325 |
| KT-TTX-GFP 0.05 mM 3MBz | Replicate 1 vs replicate 2 | 0.5969 |
| KT-TTX-GFP 0.05 mM 3MBz | Replicate 1 vs replicate 3 | 0.3664 |
| KT-TTX-GFP 0.05 mM 3MBz | Replicate 2 vs replicate 3 | 0.3127 |
| KT-TTX-GFP 0.1 mM 3MBz | Replicate 1 vs replicate 2 | 0.6501 |
| KT-TTX-GFP 0.1 mM 3MBz | Replicate 1 vs replicate 3 | 0.6154 |
| KT-TTX-GFP 0.1 mM 3MBz | Replicate 2 vs replicate 3 | 0.1598 |
| KT2440-GFP | Replicate 1 vs replicate 2 | 0.6854 |
| KT2440-GFP | Replicate 1 vs replicate 3 | 0.5568 |
| KT2440-GFP | Replicate 2 vs replicate 3 | 0.3058 |

Table S3: Kolmogorov-Smirnov test values to evaluate differences in the distribution of each cytometry replicate. Populations with a similar distribution show values KS ¡0.3, while populations with differences usually display values KS¿0.3. P-values are not shown. According to their very low magnitude they can be considered ˜ 0 for all the tests.

| Parameter | Typical value | Units | Reference |
|---|---|---|---|
| $\alpha_L$ | $4.0 \times 10^{-3}$ | $\mathrm{mM\,h^{-1}}$ | [1, 2] |
| $\alpha_R$ | $4.0 \times 10^{-3}$ | $\mathrm{mM\,h^{-1}}$ | [1, 2] |
| $\alpha_Y$ | $1\text{–}7 \times 10^{-3}$ | $\mathrm{mM\,h^{-1}}$ | [3, 2] |
| $\beta_Y$ | $1.0 \times 10^{-3}$ | $\mathrm{mM\,h^{-1}}$ | [3] |
| $\delta_L$ | 1.0 | $\mathrm{h^{-1}}$ | [4, 5] |
| $\delta_R$ | 1.0 | $\mathrm{h^{-1}}$ | [4, 5] |
| $\delta_Y$ | $2.3 \times 10^{-1}$ | $\mathrm{h^{-1}}$ | [6, 7] |
| $K_L$ | $1.0 \times 10^{-6}$ | mM | [8, 9] |
| $K_R$ | $1.0 \times 10^{-6}\text{–}1.0 \times 10^{-4}$ | mM | [10, 11] |
| $n_L$ | 1.5 | — | [12, 13] |
| $n_R$ | 1.0 | — | [14, 15] |
| $K_I$ | $1.5 \times 10^{-2}$ | mM | [16, 13] |
| $m$ | 1.0 | — | [17, 13] |

Table S4: Parameter values used in the model of the NOT gates.
